# Protection of grapevine pruning wounds against *Phaeomoniella chlamydospora* and *Diplodia seriata* by biological and chemical methods

**DOI:** 10.1101/2020.05.26.117374

**Authors:** María del Pilar Martínez-Diz, Emilia Díaz-Losada, Ángela Díaz-Fernández, Yolanda Bouzas-Cid, David Gramaje

## Abstract

The grapevine trunk diseases (GTDs) Botryosphaeria dieback and esca threaten the sustainability of the grapevine industry worldwide. This study aimed to evaluate and compare the efficacy of various liquid (pyraclostrobin + boscalid and thiophanate methyl) and paste (paste + tebuconazole) formulation fungicide treatments, and biological control agents (*Trichoderma atroviride* SC1 and *T. atroviride* I-1237), for their potential to prevent infection of grapevine pruning wounds by *Diplodia seriata* and *Phaeomoniella chlamydospora* in two field trials over two growing seasons. Treatments were applied to freshly pruned wounds following their label dosages recommendations. After 24 hours, wounds were artificially inoculated with 400 spores of *D. seriata* or 800 spores of *P. chlamydospora*. Isolations were made from the treated pruning wounds after 12 months to evaluate the efficacy of the treatments. Fungicide formulations were superior to *Trichoderma*-based treatments for the control of both pathogens during both growing seasons, with mean percent disease control of 44 to 95% for *D. seriata* and 46 to 67% for *P. chlamydospora.* Pyraclostrobin + boscalid was the most effective treatment. *Trichoderma atroviride*-based treatments did not reduce infection by *D. seriata* or *P. chlamydospora* compared to the untreated inoculated control in both vineyards and seasons. This study represents the first vineyard assessment of several chemical and biological treatments to protect pruning wounds against GTDs fungi in Europe and provides growers with tangible preventative control practices to minimize yield losses due to GTDs.

## 1 Introduction

Botryosphaeria dieback and esca are two of the most harmful grapevine trunk diseases (GTDs) affecting vineyards in all major grape-producing areas worldwide. They currently are among the main biotic threats to the economic sustainability of viticulture reducing yields, productivity and longevity of vines and vineyards (Gramaje et al., 2018). Yield losses of 30-50% have been reported by Botryosphaeria dieback in highly infected vineyards of North America (Milholland, 1991). The economic impact of Botryosphaeria dieback along with another GTD such as Eutypa dieback in California was estimated to be $USD260 million per year (Siebert, 2001). Esca incidence has reached up to 80% in several vineyards of Southern Italy (Romanazzi et al., 2009), and 12% of vineyards in France are currently no longer economically viable, due mainly to esca, with an annual estimated loss of €1 billion (Lorch, 2014).

Botryosphaeria dieback is currently associated with 26 botryosphaeriaceaous taxa in the genera *Botryosphaeria*, *Diplodia*, *Dothiorella*, *Lasiodiplodia*, *Neofusicoccum*, *Neoscytalidium*, *Phaeobotryosphaeria*, and *Spencermartinsia* (Úrbez-Torres, 2011; Pitt et al., 2013a, 2013b, 2015; Rolshausen et al., 2013; Yang et al., 2017) with the species *Diplodia seriata* being one of the most frequently isolated fungi from diseased vines in several grape growing regions such as Australia (Savocchia et al., 2007), California (Úrbez-Torres et al., 2010), Chile (Auger et al., 2004), China (Yan et al., 2013), France (Larignon et al., 2001), Mexico (Úrbez-Torres et al., 2008), Portugal (Phillips, 2002), South Africa (van Niekerk et al., 2004) and Spain (Luque et al., 2014). Botryosphaeria dieback frequently shows as complete absence of spring growth from affected spurs due to necrosis formation in wood vascular tissues with bud-break failure, and shoot and trunk dieback (Úrbez-Torres, 2011). Wood symptoms are characterized by wedge-shaped perennial cankers and dark streaking in spurs, cordons and trunks vascular tissues usually beginning in pruning wounds (Úrbez-Torres et al., 2010).

Esca is mainly caused by the fungus *Phaeomoniella chlamydospora* along with *Phaeoacremonium minimum* and other *Phaeoacremonium* spp. (Gramaje et al., 2015), some *Cadophora* spp. (Travadon et al., 2015), and several basidiomycetous taxa belonging to genera *Inocutis*, *Inonotus*, *Fomitiporella*, *Fomitiporia*, *Phellinus*, and *Stereum* (Cloete et al., 2015). The most characteristic external symptoms of the chronic esca comprise multiple banding discolourations on leaves known as ‘tiger-stripe’ pattern (Surico, 2009; Gubler et al., 2015). Internal wood symptoms involve black spots in the xylem vessels, longitudinal brown to black vascular streaking, and white to light yellow soft rot that frequently develops in wood of older vines (Fischer, 2002; Lecomte et al., 2012). Apoplectic esca form is characterized by a sudden and unexpected wilting of the whole vine or one/several arms or shoots (Lecomte et al., 2012).

Infection of grapevines by GTD fungal pathogens primarily occurs through annual pruning wounds made during the dormant season (Gramaje et al., 2018). Pycnidia of Botryosphaeriaceae spp. and *P. chlamydospora* develop from dead/cankered wood, old pruning wounds, grapevine canes, crevices, cracks and on the bark of infected grapevines (Úrbez-Torres and Gubler, 2011; Baloyi et al., 2016), and in the case of *P. chlamydospora*, mycelium on infected wood can also be a source of conidia (Edwards and Pascoe, 2001; Edwards et al., 2001; Baloyi et al., 2016). Fruiting bodies of these fungi can also be found in pruning debris left in the vineyard, thus becoming a potential inoculum source for new infections (van Niekerk et al. 2010; Úrbez-Torres, 2011; Elena and Luque, 2016b).

Conidia release of Botryosphaeriaceae spp. and *P. chlamydospora* has been shown to be primarily correlated with rain events (Larignon and Dubos, 2000; Eskalen and Gubler, 2001; Kuntzmann et al., 2009; van Niekerk et al., 2010; Úrbez-Torres et al., 2010; Valencia et al., 2015). The dynamics of *P. chlamydospora* dispersal in Spain were recently described by an epidemiological equation that integrated the effects of both rain and temperature (González-Domínguez et al., 2020). Conidia of Botryosphaeriaceae spp. has been shown to be primarily dispersed by rain splash (Úrbez-Torres et al., 2010), while inoculum of *P. chlamydospora* is predominantly aerially dispersed (Larignon and Dubos, 2000; Eskalen and Gubler, 2001; Gubler et al., 2015; Quaglia et al., 2009). Infection occurs when conidia land on exposed and susceptible pruning wounds, germinate in xylem vessels and colonize the vine spur, cordon and trunk (Mostert et al., 2006; Epstein et al., 2008; Gubler et al., 2013; Moyo et al., 2014).

Susceptibility of pruning wounds to GTD pathogens is mainly dependent on the time of pruning, and the period between pruning and possible infection case. Several studies using artificial spore inoculations showed that susceptibility of grapevine pruning wounds is high when fungal infection occurs at the moment of pruning but decreases as the period between pruning and infection increases up to several weeks or months (Petzold et al., 1981; Munkvold and Marois, 1995; Eskalen et al., 2007; Serra et al., 2008; Úrbez-Torres and Gubler, 2011), with seasonal variation reported between grape regions caused primarily by climatic differences (Gramaje et al., 2018).

Protection of pruning wounds is essential for the management of Botryosphaeria dieback and esca in grapevine, especially if adopted early in the vineyard lifespan (Kaplan et al., 2016; Sosnowski and McCarthy, 2017). The efficacy of fungicide wound treatments against Botryosphaeriaceae spp. and *P. chlamydospora* has been demonstrated in Australia (Pitt et al., 2012), California (Rolshausen et al., 2010), Chile (Díaz and Latorre, 2013), New Zealand (Amponsah et al., 2012; Sosnowski and Mundi, 2019) and South Africa (Mutawila et al., 2015). The use of physical barriers such as paints and pastes formulated with or without fungicides have also shown to be effective to control infections caused by Botryosphariaceae fungi and *P. chlamydospora* (Epstein et al., 2008; Rolshausen et al., 2010; Pitt et al., 2012; Díaz and Latorre, 2013).

The high restrictions that most effective chemical active ingredients are currently facing in Europe because of environmental and human health risks (Larignon et al., 2008; Spinosi et al., 2009), make indispensable address new alternatives for controlling GTDs. Over the last years, research on biological control of GTD fungi with antagonistic microorganisms has shown promising results primarily under controlled conditions (Alfonzo et al., 2009; Mutawila et al., 2011a; Haidar et al., 2016; Rezgui et al., 2016; Álvarez-Pérez et al., 2017; Daraignes et al., 2018; Mondello et al., 2018; Andreolli et al., 2019; Del Frari et al., 2019; Mondello et al., 2019; Trotel-Aziz et al., 2019; Niem et al., 2020). Field trials with biological control agents (BCAs) have shown variable results for preventing infection by Botryosphaeriaceae and esca fungi (Kotze et al., 2011; Mutawila et al., 2011b, 2015, 2016; Mounier et al., 2014; Reis et al., 2017; Martínez-Diz et al., 2020a).

To our knowledge, no comparative studies to evaluate the efficacy of chemical and BCA products as pruning wound protectants against GTD fungi have been performed in Europe so far. Four pruning wound treatments are currently registered in Spain for the control of GTD fungi: three *Trichoderma*-based biological products, namely Esquive, Blindar and Vintec, and Tessior, a liquid polymer containing boscalid and pyraclostrobin (MAPA, 2020). In addition, thiophanate methyl is registered in Spain against fungal trunk pathogens in almond (MAPA, 2020). The aim of this study was to evaluate and compare the efficacy of various liquid and paste formulation fungicide treatments, and BCAs, for their potential to prevent infection of grapevine pruning wounds by *D. seriata* and *P. chlamydospora* in field trials. The products assessed were those registered in Spain for control of fungal trunk pathogens or other diseases on grapevine and/or other hosts.

## 2 Materials and methods

### 2.1 Location and characteristics of the experimental vineyards

The assays were carried out at two commercial vineyards located in O Barco de Valdeorras, Galicia region (Spain), in 2018 and 2019. The vineyards were planted on 1981 (37-years-old) and 1989 (29-years-old) with ‘Godello’ cultivar grafted onto 110 Richter rootstock. Vines were spaced 120 cm from center to center, and with an interrow spacing of 225 cm, trained as bilateral cordons in a trellis system with a spur-pruning (Royat).

Vineyards were less than 500 m apart and had very similar climates. Standard cultural practices were used in both vineyards during the growing season, and the management of powdery and downy mildews was performed using only wettable sulphur and copper compounds applied at label dosages and following Integrated Pest Management (IPM) guidelines, respectively, when required. At the beginning of the study (2018), about 8% and 12% of vines had shown GTDs symptoms in each vineyard, respectively. The presence and evolution of GTDs symptoms have been inspected biannually from 2014 to present in plots of 1,500 vines at both vineyards. GTDs symptoms detected during inspection were associated mainly with esca such as tiger-pattern foliar necrosis, and shoots, arm and/or cordon death.

Both vineyards were located less than 4 km to an automatic weather station owned by MeteoGalicia (Weather Service of Galician Regional Government, Xunta de Galicia) and its climatic data was considered to be representative.

### 2.2 Fungal isolates and inoculum preparation

*Diplodia seriata* isolate CJL-398 and *Phaeomoniella chlamydospora* isolate BV-130 were used. *P. chlamydospora* BV-130 was selected due to its high virulence on grapevine in previous assays (Martínez-Diz et al., 2019). This strain was isolated from a 43-year-old esca diseased vine cultivar ‘Tempranillo’ grafted onto ‘41 Berlandieri’ rootstock in 2015*. D. seriata* JL-398 was the most virulent isolate among 14 in a detached grapevine cane assay (Elena et al., 2015a). This strain was isolated from cankers and wood necrosis of grapevine.

Conidial suspensions of each pathogen were used for artificial inoculations in the field and inoculum was obtained using methods similar to those described by Elena and Luque (2016a). In the case of *D. seriata*, a mycelial plug previously plated on Potato Dextrose Agar (PDA, Conda Laboratories, Spain) at 25°C for 7 days was cultured upside down over the center of a water agar (WA, Conda Laboratories, Spain) plate. Maritime pine (*Pinus pinaster* L.) needles were cut to 1 cm long fragments and then sterilized in an autoclave following the standard protocol of 121°C for 20 min. Then, approximately 20 sterile needles fragments were placed on the WA media surface surrounding the *D. seriata* mycelial plug at about 1 to 1.5 cm and plates were incubated under warm white fluorescent and near ultraviolet light for a 12-h photoperiod regime at 25°C for 4 weeks until pycnidia formation. The day before inoculation, pine needles fragments (about n=40) with *D. seriata* pycnidia were placed along with 30 ml of sterile distilled water (SDW) in a beaker. The solution was kept overnight (about 16 h) at 4°C in permanent agitation with the aid of a magnetic stirrer to induce conidia release from the pycnidia and prevent conidia germination. The inoculation day, the resulting solution was vacuum-filtered through a 60-mm Steriflip filter (Millipore Corporation, Billerica, MA) to get a cleaner suspension. Then, conidial suspension was adjusted to 2×10^4^ conidia mL^−1^ using a hemocytometer (Brand™ Blaubrand™ Neubauer Counting Chamber, Thermo Fisher Scientific Inc., MA, USA).

*Phaeomoniella chlamydospora* strain was grown on PDA plates at 25°C for 3 weeks. Same day of inoculation, conidia were released from cultures by adding 10 ml of SDW and gently scraping with a sterile stick and the collected suspension adjusted to a concentration of 4×10^4^ conidia mL^−1^ based on counts from the hemocytometer. Both conidial suspensions were stored at 4°C until inoculation time to avoid early conidia germination.

Spore germination was assessed for both fungal trunk pathogens by placing four drops of the spore suspension on a PDA plate, which was then incubated at 25°C under fluorescent light for a 12-h photoperiod. After approximately 24 h, a glass cover slip was placed over each drop area on the PDA. The number of non-germinated spores over a total of 100 in each drop was counted using an optical microscope (Nikon Eclipse E400) at 100x magnification. The mean percentage of germinated spores was determined.

### 2.3 Pruning wound protection treatments

Wound protection treatments tested in the present assay are listed in Table 1. We evaluated the efficacy of two chemical and two BCA formulated products, and also a paste mixed with a fungicide. In general, the chemical and biological products assessed were those commercially formulated and currently registered and available in Spain for control of fungal trunk pathogens, except tebuconazole (Song), which is registered to control botrytis bunch rot (*Botrytis cinerea*) and powdery mildew (*Erysiphe necator*) in grapevine. Applications rates were selected based on the registered label dosages recommendations. Liquid formulations were prepared by the suspension of the products in tap water, which is the procedure normally used for spraying vineyard treatments in Galicia region. Pyraclostrobin + boscalid (Tessior) treatment contains a liquid polymer and it is already formulated to be directly sprayed to pruning wounds without any previous mixing. Paste treatment was prepared by mixing a liter of the paste formulation (Master) with 80 ml of tebuconazole (Song).

**Table 1.**
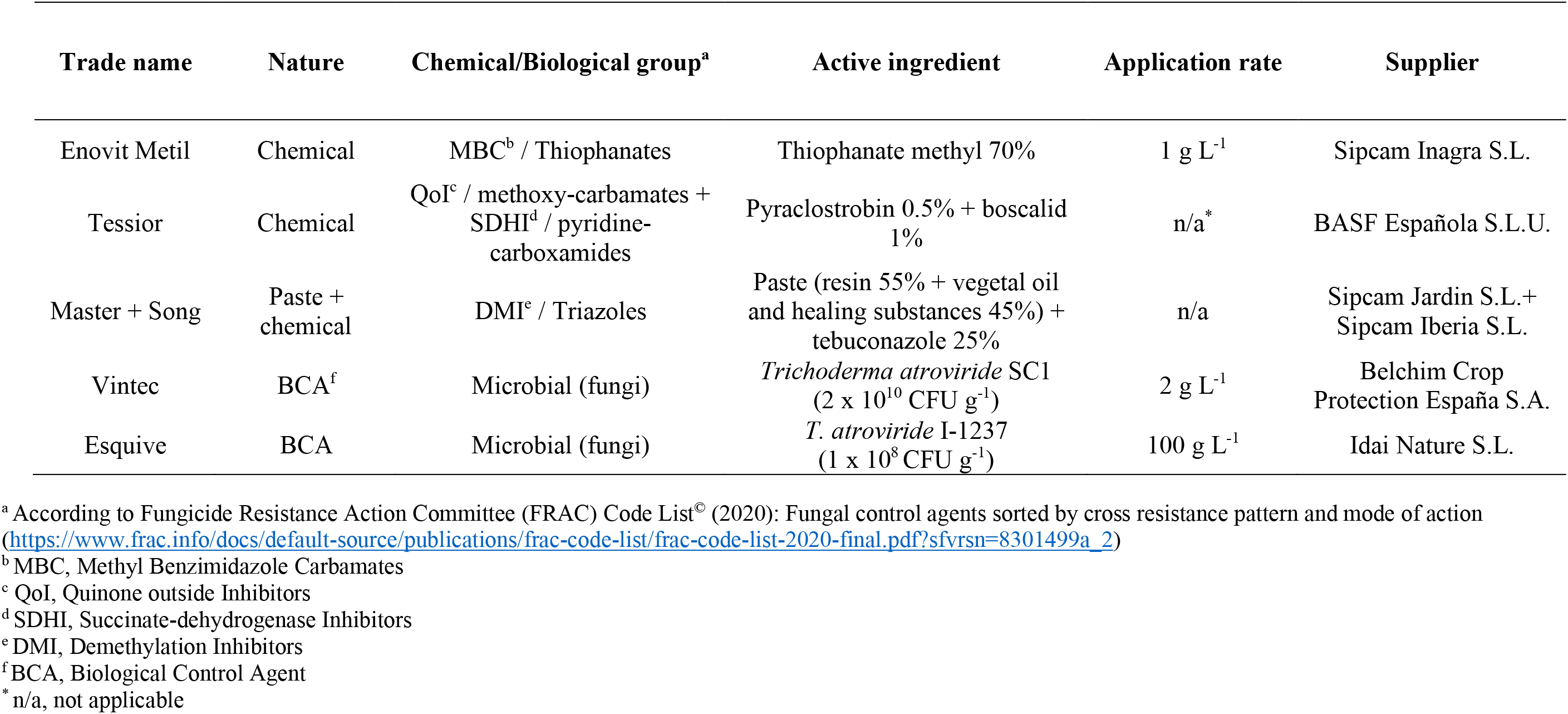
Pruning wound treatments evaluated for control of *Diplodia seriata* and *Phaeomoniella chlamydospora* under field conditions.

Regarding BCA treatments, the conidia viability of both *Trichoderma atroviride* strains (SC1 and I-1237) in the commercial products was tested to be at a minimum of 85% before the assay was set up (Pertot et al., 2016). A serial dilution of the conidia suspension was plated on PDA and the colony-forming units were counted after 24-48 h incubation at room temperature.

### 2.4 Field assay and experimental design

On 19 February 2018, 1-year-old canes of all vines to be treated were spur-pruned to three buds using secateurs in both vineyards, coinciding with the common pruning time in this region of Spain. Wounds treatments were applied by hand until runoff within 2 h after pruning to three wounds per vine. Liquid formulations were applied using a 500 ml hand-held spray bottle with a plastic shield on the nozzle to minimise spray drift and the paste formulation were applied with the aid of a paintbrush. Untreated controls, positive (artificially inoculated, IC) and negative (non-artificially inoculated, NC) were mock treated with sterile distilled water (SDW).

On the following day, wounds were moistened by spraying with SDW immediately prior to inoculation with the fungal trunk pathogens and a drop of Tween 20 (Sigma-Aldrich, San Luis, MO, USA) was added to each conidial suspension as a surfactant to assist spreading the spores over the pruning wound surface (Sosnowski and Mundi, 2019). Approximately 400 and 800 conidia of *D. seriata* and *P. chlamydospora*, respectively, suspended in a drop of 20 μl of SDW were then applied per wound using a micropipette. All pruning wounds were inoculated with the pathogen inoculum except NC controls, which were mock inoculated with a drop of 20 μl of SDW alone instead and being exposed to natural infection. Inoculum drops placed onto the pruning wounds were left to air dry (from some minutes to 1 h) before being wrapped with Parafilm M (Pechiney Plastic Packaging, Chicago, IL, USA) to avoid fast dehydration and favour fungal spores’ penetration into xylem vessels. Due care was taken to avoid the rain for the entire duration of the trials set up, namely pruning, wound treatments application and artificial fungal inoculation (2 days).

The experiment was set up as a randomized block design with three replicates of ten plants (thirty canes) per wound protectant treatment and pathogen in each vineyard. Three replicates of ten plants per pathogen were also used for IC in each vineyard. Additionally, three replicates of ten plants were used as NC in each vineyard. The experiment was repeated the following season (2019–20), with pruning and wound treatments applied on 12 February 2019, and artificial fungal inoculations on 13 February 2019.

### 2.5 Fungal recovery and identification

Canes were harvested from vines above the second bud (about 10 cm long pieces) approximately 12 months after artificial inoculation and stored in a 4°C cool room prior to laboratory assessment. Bark was first removed using a sharp knife from each cane. Then, canes were surface sterilised for 1 min in 33% sodium hypochlorite (commercial 40 g Cl/l) and rinsed twice for 1 min each in SDW. After air drying on sterile filter paper to remove moisture excess, each cane was cut into small pieces (about 12 mm^2^) taken from the margin between discoloured or dead and live or apparently healthy wood tissue using sterilised secateurs. Five wood fragments were plated onto each of two plates of Malt Extract Agar (MEA) amended with 0.35 g l^−1^ of streptomycin sulphate (Sigma-Aldrich, St. Louis, MO, USA) (MEAS) giving a total of ten wood pieces per cane. Cultures were incubated at 25°C under warm fluorescent light for a 12-h photoperiod and inspected daily for 15 days. All growing fungal colonies were transferred to PDA plates and then assessed for the presence or absence fungal mycelial growth resembling *D. seriata*, *P. chlamydospora* or *Trichoderma* spp.

Identification of GTD fungal cultures was then assessed under a stereoscopic (Olympus SZX9, Olympus Corporation, Tokyo, Japan) and optical microscopes (Nikon Eclipse E400, Nikon Corporation, Tokyo, Japan) based on cultural and morphological features previously described including colony growth pattern, colour, mycelial and other characteristics such as conidial shape, size and colour (Crous and Gams, 2000; Phillips et al., 2007). Identity of GTD fungal isolates and *Trichoderma* spp. was confirmed by molecular methods. Fungal DNA was extracted from fresh mycelium after 3 weeks of incubation in PDA using the E.Z.N.A. Plant Miniprep Kit (Omega Bio-Tek, Doraville, GA, USA) following manufacturer’s instructions. *D. seriata* was confirmed by sequencing part of the translation elongation factor 1-using the primer pairs EF1F-EF2R (Jacobs et al., 2004). *P. chlamydospora* was detected by PCR using the primers Pch1-Pch2 (Tegli et al., 2000). Identity of *Trichoderma* spp. was confirmed at species level by sequencing the ITS region using the universal primers ITS1F/ITS4 (Gardes and Bruns, 1993). All PCR products were visualized in 1% agarose gels (agarose D-1 Low EEO, Conda Laboratories) and sequenced in both direction by Eurofins GATC Biotech (Cologne, Germany).

### 2.6 Data analysis

Efficacy of each wound treatment was calculated as mean percentage recovery (MPR) of *D. seriata* and *P. chlamydospora* from each cane per treatment (Sosnowski et al., 2008, 2013). Data were checked for normality and homogeneity of variances prior to statistical analyses and transformed when required into the arcsine of the square root of the proportion (MPR/100)^1/2^. The statistical analysis of the experimental results was carried out in a two-way ANOVA with blocks and treatments as independent variables, and MPR (%) as dependent variable. Mean percentage disease control (MPDC) was also determined as the reduction in MPR (%) as a proportion of the artificially inoculated control (IC) (MPDC=100 × [1 – (MPR treatment/MPR IC)]) (Sosnowski et al., 2008, 2013). Means were compared with ICs by the Student’s *t* least significant difference (LSD) at *P* < 0.05. Data from all experiments were analysed using the Statistix 10 software (Analytical Software).

## 3 Results

### 3.1 Wound treatment evaluation against Diplodia seriata

During 2018-19 and 2019-20 seasons, *D. seriata* spore germination on PDA was 94% and 98.5%, respectively, and it was recovered from 58 and 68% of IC wounds, respectively (Table 2). *D. seriata* was recovered from 1% of NCs wounds at both seasons. Analysis of variance showed that there were significant differences in the relative recovery data from the different treatments between seasons (*P*<0.05). No significant differences were found in the recovery data between vineyards in each season (2018-19, *P*=0.904; 2019-20, *P*=0.593), so data from each vineyard were combined and the analysis was performed separately for each season (Table 2).

**Table 2.**
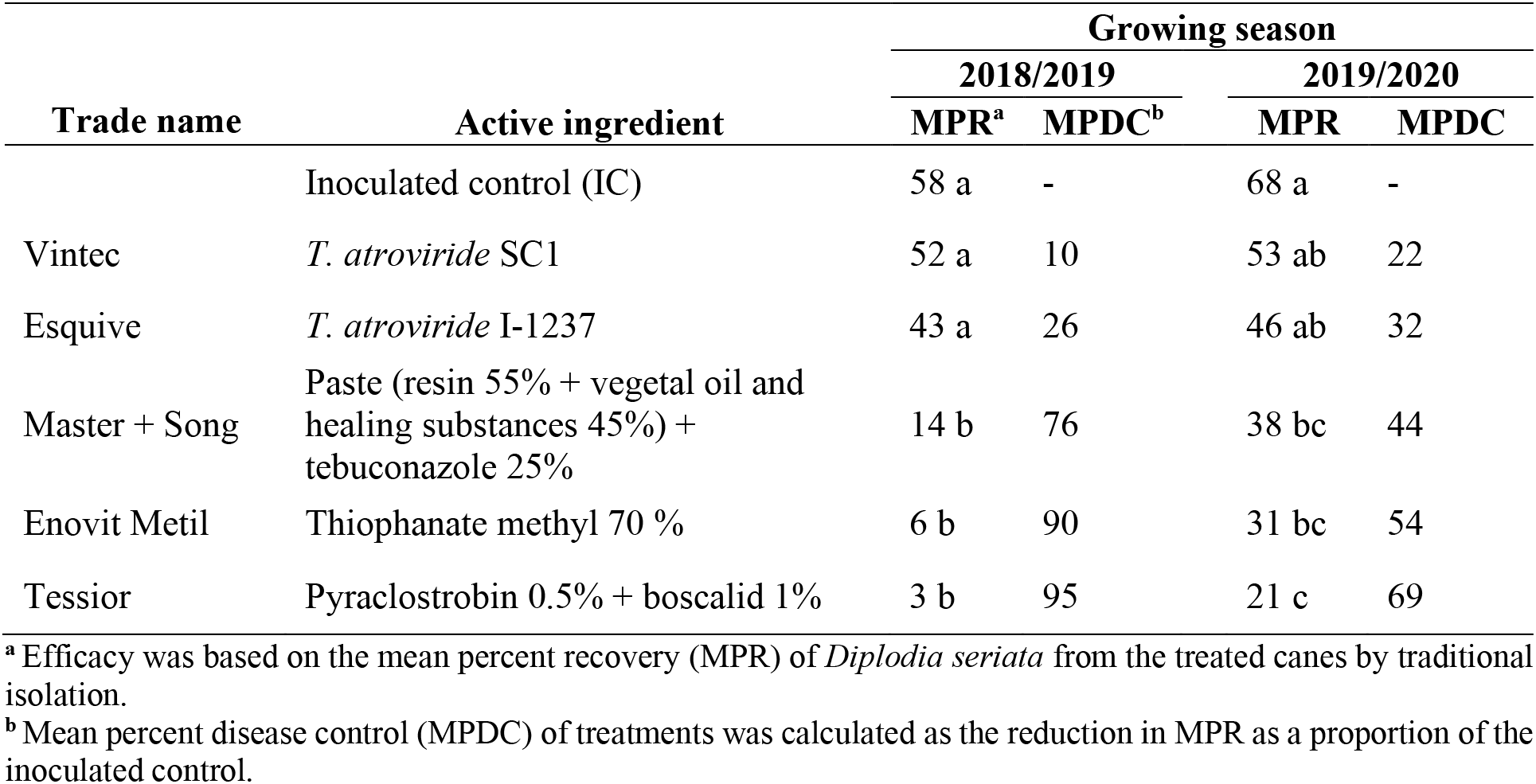
Efficacy of wound treatments when applied 24 h before inoculation with *Diplodia seriata* in two growing seasons.

| Trade name | Active ingredient | Growing season |  |  |  |
| --- | --- | --- | --- | --- | --- |
|  |  | 2018/2019 |  | 2019/2020 |  |
|  |  | MPR<sup>a</sup> | MPDC<sup>b</sup> | MPR | MPDC |
|  | Inoculated control (IC) | 58 a | - | 68 a | - |
| Vintec | *T. atroviride* SC1 | 52 a | 10 | 53 ab | 22 |
| Esquive | *T. atroviride* I-1237 | 43 a | 26 | 46 ab | 32 |
| Master + Song | Paste (resin 55% + vegetal oil and healing substances 45%) + tebuconazole 25% | 14 b | 76 | 38 bc | 44 |
| Enovit Metil | Thiophanate methyl 70 % | 6 b | 90 | 31 bc | 54 |
| Tessior | Pyraclostrobin 0.5% + boscalid 1% | 3 b | 95 | 21 c | 69 |

<sup>a</sup> Efficacy was based on the mean percent recovery (MPR) of *Diplodia seriata* from the treated canes by traditional isolation.
<sup>b</sup> Mean percent disease control (MPDC) of treatments was calculated as the reduction in MPR as a proportion of the inoculated control.

Treatment with pyraclostrobin + boscalid, thiophanate methyl, and the paste + tebuconaloze significantly reduced the MPR of *D. seriata* from pruning wounds with respect to the IC at both seasons (*P*<0.05) (Table 2). During 2018-19 season, pyraclostrobin + boscalid, thiophanate methyl, and the paste + tebuconaloze provided MPDC of 95, 90 and 76%, respectively, whereas these products provided MPDC of 69, 54 and 44%, respectively, during 2019-20 season. During both seasons, there was not a significant treatment effect with *Trichoderma*-based wound protectants (*P*>0.05). During 2018-19 season, *T. atroviride* SC1 and *T. atroviride* I-1237 provided MPDC of 10, and 26%, respectively, while these products provided MPDC of 22 and 32%, respectively, during 2019-20 season.

### 3.2 Wound treatment evaluation against Phaeomoniella chlamydospora

During 2018-19 and 2019-20 seasons, *P. chlamydospora* spore germination on PDA was 89% and 93%, respectively, and it was recovered from 27 and 42% of IC wounds, respectively (Table 3). *P. chlamydospora* was recovered from 0% and 3% of NCs wounds during 2018-19 and 2019-20 seasons, respectively. There were no significant differences in the relative recovery data from the different treatments between vineyards (*P*=0.500) and seasons (*P*=0.080), so data were combined for analysis.

**Table 3.** Efficacy of wound treatments when applied 24 h before inoculation with *Phaeomoniella chlamydospora*.

| Trade name | Active ingredient | MPR<sup>a</sup> | MPDC<sup>b</sup> |
| --- | --- | --- | --- |
|  | Inoculated control (IC) | 36 ab | - |
| Vintec | *T. atroviride* SC1 | 45 a | 0 |
| Esquive | *T. atroviride* I-1237 | 30 ab | 17 |
| Enovit Metil | Thiophanate methyl 70% | 19 bc | 46 |
| Tessior | Pyraclostrobin 0.5% + boscalid 1% | 18 c | 51 |
| Master + Song | Paste (resin 55% + vegetal oil and healing substances 45%) + tebuconazole 25% | 12 c | 67 |

<sup>a</sup> Efficacy was based on the mean percent recovery (MPR) of *Phaeomoniella chlamydospora* from the treated canes by traditional isolation.
<sup>b</sup> Mean percent disease control (MPDC) of treatments was calculated as the reduction in MPR as a proportion of the inoculated control.

There was a significant treatment effect (*P*<0.05) with the paste + tebuconaloze and pyraclostrobin + boscalid treatments reducing MPR of *P. chlamydospora* to 12 and 18% compared with 36% from the IC wounds (MPDC of 67 and 51%; Table 3). There was not a significant treatment effect with thiophanate methyl and *Trichoderma*-based wound protectants (*P*>0.05). Thiophanate methyl provided MPDC of 46%, whereas *T. atroviride* SC1 and *T. atroviride* I-1237 provided MPDC of 0 and 17%, respectively.

### 3.3 Trichoderma-based treatments colonization

The conidia viability was on average of 97% and 95% for *T. atroviride* SC1 during 2018-19 and 2019-20 seasons, respectively. Regarding *T. atroviride* I-1237, the conidia viability was 94% during 2018-19 season and 96% during 2019-20 season. *Trichoderma* spp. were exclusively recovered from pruning wounds treated with *Trichoderma*-based formulations at varying levels. There were no significant differences in the relative recovery data between vineyards (*P*=0.180) and seasons (*P*=0.075). During 2018-19, recovery percentages were 5 and 10% for *T. atroviride* SC1 and *T. atroviride* I-1237, respectively. During 2019-20, recovery percentages were 9% and 14% for *T. atroviride* SC1 and *T. atroviride* I-1237, respectively.

### 3.4 Weather data

During 2018-19 season, the average of the daily mean temperature and relative humidity in the week from the day of pruning and wound treatments application (from 19 to 25 February 2018) was 6.5°C and 67.5%, respectively, with no rain events in that period. The average of daily mean temperature, daily mean relative humidity and accumulated rainfall for the whole month of February 2018 was 5.8°C, 76.6% and 84.2 mm, respectively, with nine rain events (of >1 mm) in total.

During 2019-20 season, the week from the day of pruning and application of wound treatments (from 12 to 18 February 2019) registered an average of the daily mean temperature of 8.2°C and a 71.1% on average of daily relative humidity. For the same period, there was only one rain event (18 February 2019) with a total rainfall of 10.6 mm. Regarding the whole February 2019 month, the average of the daily temperature was 8.2°C and of the daily relative humidity 73.3%. The total rainfall in the same month was 37 mm received in a total of four rain events.

## 4 Discussion

The present study represents the first vineyard comparison of the efficacy of paste and liquid fungicides, and BCA treatments to protect pruning wounds against GTDs fungi in Europe. Considering the high incidence of GTDs, particularly esca, and the restrictions on the use of chemicals in Europe (Mondello et al., 2018), this study provides growers with tangible preventative control practices to minimize yield losses due to GTDs. By focussing on products already registered for control of trunk diseases in almond or foliar diseases of grapevines in Spain, the lower cost of label extension compared to new product registration will increase the likelihood and success of registration for GTDs. *D. seriata* was chosen to represent Botryosphaeria dieback, because it is one the most common cited Botryosphaeriaceae species occurring on grapevines worldwide and is reported to be a virulent species in Spain (Luque et al., 2009; Elena et al., 2015b). *P. chlamydospora* was chosen to represent esca, because it is the most frequently isolated species from affected vines in most grape growing regions worldwide (Berstch et al., 2013; Gubler et al., 2015).

Our results demonstrate that paste and liquid fungicide formulations were superior to *Trichoderma*-based treatments for the control of *D. seriata* and *P. chlamydospora.* All paste and liquid fungicide treatments tested reduced recovery of both pathogens from inoculated wounds compared with the untreated inoculated control, with the exception of thiophanate methyl for *P. chlamydospora*. Similar results were observed in other studies where several fungicides and BCA treatments were compared as pruning wound protectants in the same field trial. In Australian vineyards, liquid and paste fungicide formulations were more effective than *Trichoderma*- and *Bacillus subtilis*-based formulations against *D. seriata* and *Diplodia mutila* (Pitt et al., 2012), and *Eutypa lata* (Ayres et al., 2017) infections, respectively. Halleen et al. (2010) also reported that fungicides were more effective than *Trichoderma* spp. against *E. lata* infection in field trials carried out in South Africa, in spite of the efficacy of *Trichoderma* treatments in reducing GTD fungal infection.

Application of pyraclostrobin + boscalid to pruning wounds provided high mean percentage of disease control (MPDC) for both pathogens. To date, only preliminary studies have been carried out in field trials in Germany (Kühn et al., 2017; Lengyel et al., 2019), Greece (Kühn et al., 2017; Samaras et al., 2019) and Spain (Kühn et al., 2017), where pyraclostrobin and boscalid (Tessior) was effective as pruning wound protectant reducing the grapevine wood infection caused by *Diplodia* spp. and *P. chlamydospora*. The application of a similar commercial product based on pyraclostrobin and boscalid without the liquid polymer (BASF516, BASF Australia Ltd, Sidney, New South Wales, Australia) showed a low efficacy against *E. lata* artificial pruning wound inoculations in Australian vineyards (Sosnowski et al., 2008). Wound applications of pyraclostrobin alone were effective for the control of *D. seriata* and *P. chlamydospora* in Chile (Díaz and Latorre, 2013) and California (Rolshausen et al., 2010) vineyards. Moreover, this active ingredient significantly reduced infections caused by fungi associated with Botryosphaeria dieback (Rolshausen et al., 2010), Eutypa dieback (Sosnowski et al., 2008, 2013; Rolshausen et al., 2010; Ayres et al., 2017), and esca (Rolshausen et al., 2010), under field conditions.

The only treatment to provide a similar level of control than pyraclostrobin + boscalid for both pathogens was the paste with tebuconazole. Accordingly, applications of paste and liquid formulations containing tebuconazole on pruning wounds of ‘Cabernet Sauvignon’ vines significantly reduced the mean vascular discolouration length and the reisolation percentage of *D. seriata* and *P. chlamydospora* in Chilean vineyards (Díaz and Latorre, 2013). In Australia, a gel and a paint with tebuconazole applied by paintbrush to freshly pruned canes reduced *E. lata* infections to 100% and 94%, respectively (Sosnowski et al., 2013). Pitt et al. (2012) also demonstrated that a tebuconazole paste formulation provided a 38% control of *D. mutila* in a trial performed in Australia. Other physical barriers containing a paste with fungicides have resulted effective at reducing pruning wound infections by other GTD fungi (Rolshausen and Gubler, 2005; Sosnowski et al., 2008; Rolshausen et al., 2010; Pitt et al., 2012). Liquid spray applications of tebuconazole were also significantly effective reducing the recovery of *D. seriata* in Australia (Pitt et al., 2012).

Thiophanate methyl was effective in reducing infection by *D. seriata*, while no significant effect was observed against *P. chlamydospora*. Similar findings were reported by Rolshausen et al. (2010) in California, where pruning wounds applications of thiophanate methyl reached a disease control of 80% for *D. seriata* infections but did not perform as well against *P. chlamydospora* with only a 52% of disease control. In Chile, Díaz and Latorre (2013) reported the efficacy of both liquid and paste formulations of thiophanate methyl to control *D. seriata* and *P. chlamydospora* infections in pruning wounds. This chemical compound was also effective in reducing the pruning wound infections caused by *P. chlamydospora* and *Neofusicoccum luteum* in field trials carried out in South Africa (Mutawila et al., 2015) and New Zealand (Amponsah et al., 2012), respectively.

Pastes and paints are considered the most reliable protectants of pruning wounds against GTD fungi, especially when they are mixed with fungicides (Moller et al., 1977; Rolshausen and Gubler, 2005; Rolshausen et al., 2010; Sosnowski et al., 2008, 2013; Díaz and Latorre, 2013). They provide a physical barrier to protect pruning wounds from GTD fungal infection while the fungicide can also act on the pathogens if the physical barrier is compromised by rain, sap flow, or cracking when drying (Sosnowski et al., 2008). However, some other studies reported no differences in effectiveness between application of acrylic paint with or without fungicides (Sosnowski et al., 2008; Mayet and Lecomte, 2014). Pastes and paints are usually applied by hand with a paint brush, unless the product contains a liquid polymer to act as a physical barrier, which is the case of Tessior commercial product. It should be noted that application by hand is more time-consuming and can be at least two to four times the application cost with a tractor mounted sprayer (Sosnowski and McCarthy, 2017). Further research is therefore required to determine the protective mechanisms of each component and if their efficacy is also influenced by other factors such as wound size, application time, and weather variables.

Species of the fungal genus *Trichoderma* have been the most investigated BCA to act as pruning wound protectant against GTDs pathogens (John et al., 2005; Halleen et al., 2010; Kotze et al., 2011; Mutawila et al., 2015, 2016; Reis et al., 2017). Our results shown that *Trichoderma atroviride*-based treatments did not reduce infection by *D. seriata* or *P. chlamydospora* compared to the untreated inoculated control in both vineyards and seasons. This is the first report to assess the efficacy of *T. atroviride* SC1 in protecting grapevine pruning wounds from infection by GTD fungi in mature vineyards. In recent research, Berbegal et al. (2020) applied *T. atroviride* SC1 to pruning wounds of 3-year-old vines but its efficacy as wound protectant against GTD pathogens was not tested in this specific plant part. In nurseries, *Trichoderma atroviride* SC1 showed high efficacy to reduce artificial (Pertot et al., 2016) or natural (Berbegal et al., 2020) *P. chlamydospora* infection when applied at different propagation stages. Preliminary results showed the efficacy of *T. atroviride* I-1237 to reduce the disease incidence and severity of *P. chlamydospora* and *N. parvum* on pruning wounds in Portuguese vineyards (Reis et al., 2017). Similarly, Mounier et al. (2014) demonstrated that spraying pruning wounds with *T. atroviride* I-1237 over two years significantly reduced the esca, and Botryosphaeria and Eutypa diebacks foliar symptoms expression, and the plant mortality rate due to GTDs in French vineyards. Dipping young grapevine plants in *T. atroviride* I-1237 during the nursery propagation process decreased *D. seriata* and *P. chlamydospora* DNA and necrotic lesion length compared to the untreated plants (Mounier et al., 2014). Other *Trichoderma* strains or *Trichoderma*-based commercial products have shown high efficacy in reducing the recovery of GTDs fungal pathogens from artificially inoculated pruning wounds under field conditions (John et al., 2005; Halleen et al., 2010; Kotze et al., 2011; Pitt et al., 2012; Mutawila et al., 2015).

These inconsistences found in the *Trichoderma* products performance among our study and previous reports could be due to different reasons. Although *Trichoderma* spp. have the ability to provide long-term protection to pruning wounds and thus preventing fungal trunk pathogens infections, they firstly need to establish itself, grow and colonize wounds instead of a simply temporal establishment (Munkvold and Marois, 1993; Mutawila et al., 2011a, 2011b). In this study, pruning wound colonization by both strains of *T. atroviride* was very low at both vineyards and seasons ranging from 5 to 14%. Environmental conditions such as the temperature at the time of application might have a negative influence in the persistence and implementation of *Trichoderma* spp. (Elmer and Reglinski, 2006; Pertot et al., 2017). According to the label recommendations of each product, *T. atroviride* SC1 formulation should be applied when environmental temperature is equal or higher than 10°C for a minimum of five hours on the day of application in the field, while *T. atroviride* I-1237 formulation is supposed to be biologically active at temperatures above 5°C. The average of the daily mean temperature experienced in the week of the trials set up was 6.5°C and 8.2°C during 2018-19 and 2019-20 seasons, respectively, a fact that could explain the low colonization performed by *Trichoderma*-based formulations. In addition, a slightly higher *Trichoderma* recovery was registered during 2019-20 season (9.4 to 13.5%) than 2018-19 (4.8 to 9.8%) probably also explained by the warmer temperatures registered in this season.

Timing of pruning within the dormant season should be adjusted to periods with mild and favourable temperature values that might lead to a better implantation and development of the BCA on pruning wounds and thus increasing its effectiveness against GTD pathogens. *Trichoderma* spp. application after pruning in late winter or early spring would likely provide higher disease control than normal pruning in winter. However, late pruning is not feasible in all vineyards. In those vineyards with limited labour force, growers need to begin pruning early in the winter to ensure completion of the activity before bud break. An alternative would be to prune in late autumn or early winter. Recent research carried out in the same grape-growing region of the present study reported low abundances of GTDs pathogens infecting naturally pruning wounds after an early pruning made in November (Martínez-Diz et al., 2020b). The age and physiological state of the vine as well as the dose and product formulation have also been suggested as factors that could have an influence on effective colonization by *Trichoderma* spp. (Schubert et al., 2008; Halleen et al., 2010; Mutawila et al., 2016). However, further research is required to confirm these hypotheses.

Label instructions of most fungicide and BCA commercial formulations to protect pruning wounds recommend their application shortly after pruning to minimize the chances of GTDs infection. In our study, we followed the official method of European countries to evaluate pruning wounds protection products against *E. lata,* which suggests carrying out fungal inoculations 24 hours after the application of preventive treatments (EPPO, 2017). Accordingly, in most of the previous pruning wound protection trials, the time elapsed between pruning wound protection and GTD fungal inoculation was 24 hours (John et al., 2005; Sosnowski et al., 2008, 2013; Halleen et al., 2010; Rolshausen et al., 2010; Kotze et al., 2011; Amponsah et al., 2012; Pitt et al., 2012; Díaz and Latorre, 2013; Ayres et al., 2017; Sosnowski and Mundi, 2019). This short time between BCA treatments application and artificial fungal inoculations could also have explained the poor performance exhibited by *Trichoderma*-based commercial formulations in our study. Previous research reported a greater biocontrol efficacy when artificial GTD pathogen infection was delayed 7 (Kotze et al., 2011; Mutawila et al., 2015) and 14 (Munkvold and Marois, 1993; John et al., 2005) days after application of *Trichoderma* spp. on pruning wounds. These findings suggest BCAs might need a period to colonise the pruning wound surface and grapevine wood to be effective, as reported by John et al. (2005).

The use of pathogen artificial inoculations is very common in the assessment of the efficacy of pruning wound protectants against GTDs to guarantee a substantial establishment of infection in untreated inoculated controls for statistical analysis (Halleen et al., 2010; Rolshausen et al., 2010; Sosnowski et al., 2008, 2013; Amponsah et al., 2012; Pitt et al., 2012; Ayres et al., 2017; Sosnowski and Mundi, 2019). In the present study, artificial inoculations with 400 (*D. seriata*) and 800 (*P. chlamydospora*) conidia were applied per wound to obtain optimal recovery percentages for robust evaluation of treatments according to doses recommendations made by Elena et al. (2015b). This fact represents a significantly higher ‘disease pressure’ than that which might be expected to occur under natural conditions. Wounds were infected naturally up to 1% by *D. seriata* and 3% by *P. chlamydospora*, in contrast with the artificially inoculated controls recovery with up to 68% and 42%, respectively. This indicates that wound protectants that showed lower efficacy rates in this study, such as BCA formulations, will most likely provide better control of both *D. seriata* and *P. chlamydospora* under ‘natural disease pressure’ in the vineyard. The efficacy of pruning wound protectants under lower artificial GTD inoculum levels or natural infections in the vineyard should be tested in future studies. Different affinities of *T. atroviride* strains for specific grapevine cultivars has been previously reported in South Africa (Mutawila et al., 2011b), and this should not be discarded as a possible cause of the low *Trichoderma* colonization rates obtained in this study.

To conclude, this study highlighted the efficacy of several fungicides with or without a physical barrier to protect grapevine pruning wounds against *D. seriata* and *P. chlamydospora* infections under field conditions. In particular pyraclostrobin + boscalid (Tessior), a registered product against GTD fungi in several countries in Europe, is recommended as pruning wound protectant to prevent infection by the most prevalent pathogens associated with Botryosphaeria dieback and esca. *Trichoderma-*based treatments showed lower efficacy against GTD fungi than that provided by fungicides and their performance seems to be related to environmental conditions and wound colonisation prior to infection by the pathogens. Good pruning practices along with strict sanitation procedures and pruning wound protection by the application of authorized products can significantly reduce the impact of GTD pathogens infections and thus increasing the lifespan of vineyards.

## Acknowledgments

This work has been funded by European funds in collaboration with the ‘Ministerio de Agricultura, Pesca y Alimentación (MAPA)’ (Spanish Government) and the ‘Xunta de Galicia’ (Regional Government) through the project ‘EVID: Innovative practices to combat grapevine diseases’ (FEADER 2017/003B). The authors acknowledge Bodegas Godeval for the provision of field sites and technical assistance for this study. We also thank Dr. J. Luque for the provision of *Diplodia seriata* isolate JL-398. M.P. Martínez-Diz was supported by the FPI-INIA program from the INIA (CPD2015-0188). D. Gramaje was supported by the Ramón y Cajal program, Spanish Government (RYC-2017-23098).

